# Extracting anomalous diffusion parameters from multi-state ensembles of short single molecule trajectories

**DOI:** 10.64898/2026.05.30.729014

**Authors:** Alisha Budhathoki, Ganesh Pandey, Jonah Galeota-Sprung, Jan-Hendrik Spille

## Abstract

Single-molecule tracking measures the stochastic motion of individual biomolecules in the cellular environment. Statistical analysis of trajectory ensembles is required to gain insight into the biophysical nature of mobility states and molecular interactions that they reflect. Mobility states can be parameterized by a generalized diffusion coefficient and anomalous exponent. Experimental constraints such as finite track length and localization precision limit how accurately these parameters can be determined. We compare the performance of analysis methods to recover the input parameters from ensembles of simulated single molecule tracks from different states spanning the range of anomalous diffusive behaviors observed in the cell nucleus. We further develop a framework to quantify error rates in the assignment of mobility states to individual molecules based on recall rates and precision. Our analysis shows that single-track analysis methods are superior to bulk methods in their ability to recover parametric descriptors from mixed populations. The most complete description is obtained by combining outputs from different tools. Our work provides a guide to assess the accuracy of analyses and obtain the most accurate parametric description of experimental single particle tracking data.

**Statement of significance:** Experimental single particle tracking data provides rich insight into molecular interactions directly in living cells. But data analysis depends critically on choosing the correct diffusion model and appropriate tools to extract accurate information. Importantly, it is usually not obvious from the output of a method whether the results are accurate or not. In this work, we use ensembles of tracks simulated with fractional Brownian motion methods to characterize the impact of track length and localization precision on analysis outcomes. We elaborate on specific strengths and weaknesses of commonly used and newly developed analysis tools to provide a template for thorough assessment and quantification of error rates in experimental data analysis.

## Introduction

Single-particle tracking (SPT) is a powerful tool to investigate how biomolecules explore the cellular environment. It has become widely adopted for studying intranuclear protein dynamics. The ease of CRISPR/Cas9 genome editing (1) for tagging of endogenous proteins, availability of bright, photo-stable ligands (2–4), as well as improved instrumentation have contributed to the continuing success of this method. SPT can reveal diffusion parameters that characterize molecular mobility (5) and target search mechanisms (6–8), binding times due e.g. to chromatin interactions (9), and drivers of structural chromatin organization (10).

To derive biophysical models of molecular exploration and target search, it is crucial to extract accurate diffusive parameters and state distributions. The majority of biomolecules will exhibit different mobility states that reflect functional interactions, e.g. transcription factors often switch between different diffusive exploration modes and also exhibit at least one bound state (6,11). These mobility states are characterized by their relative abundance and parametric descriptors, i.e. the generalized diffusion coefficient and an anomalous exponent that reflect spatial exploration behavior. Additional readouts can include residence times in, and transition rates between mobility states (12,13). The most accurate results in particular for the characterization of individual molecules will be obtained from long observation times, i.e. higher numbers of samples of a molecule in a given state, but practically, behavior of the molecules themselves often limits track length (14). Approaches to extend trajectory length require specialized sample labeling (15) or instrumentation strategies (16) that may not be universally applicable. It is therefore important to understand the requirements and limitations of analysis tools to extract the most accurate descriptors within the constraints of experimental realities.

All analysis methods have in common that they assume a biophysical diffusion model to determine these parametric descriptions. Model parameters are optimized to achieve the best fit of the model to experimental data distributions. The interpretation of single particle mobility thus depends on the choice of model. Not all methods are suited to accommodate all models.

Choosing an incorrect model leads to misinterpretation of the results, e.g. when a free Brownian motion model is used to parameterize sub-diffusive motion in jump distance analysis, or when single state mean-square displacement analysis is applied to an ensemble of mobility states. It can be difficult to decide based on the output of a given tool whether the chosen model was indeed appropriate. Neither goodness of fit parameters nor inspection of fit residuals are sufficient to determine if a model is appropriate. Incorrect models can yield satisfying fits, particularly in the analysis of mixed ensembles where an unknown number of mobility states is required.

A ground truth reference typically does not exist for experimental data. Here, we use fractional Brownian motion simulation (17) to generate synthetic ensembles of mixed anomalous diffusion states with known parameters to benchmark the performance of commonly used SPT analysis tools. We assess their ability to report relative fractions, generalized diffusion coefficients, and anomalous exponents. To this end, we simulate anomalous diffusion ensembles. We determine how track length and localization precision affect the outcomes.

Beyond general parametric descriptions of mobility states, it is of interest to determine which state a given single particle observation represents. This information is of particular interest when trajectories can be linked to events or spatial landmarks to determine if an association correlates with changes in molecular mobility. We quantify recall and precision of two different analysis methods that allow us to assign individual tracks to one of the subpopulations in a mixed ensemble. We find that single-track methods are hampered by short track length more so than localization error. Nevertheless, we find that single track analysis is best suited for recovering diffusive parameters of sub-populations.

This investigation was prompted by recent experiments in which we acquired single-molecule tracks of Rpb1, the largest subunit of the RNA Polymerase II complex, in mouse embryonic stem cell nuclei, and specifically within transcription condensates (18). The presence of multiple mobility states in the ensemble of tracks, previously reported by others (19–21), raised the question how accurate the reported diffusion parameters are, and whether it is possible to reliably identify which state an individual molecule is in.

In this manuscript, we generate an *in silico* mimic of the experimental dataset with a known ground truth. We compare classical bulk analysis methods, mean squared displacement (MSD) and jump distance distribution (JDD) analysis (22), as well as novel approaches based on the characterization of individual tracks, single-track MSD (stMSD)(23) followed by Gaussian mixture modeling (GMM) (18), and state array single particle tracking (saSPT) (17) in their ability to report diffusion parameters and relative fractions of sub-populations as well as their accuracy in determining the state of individual tracks.

## Materials and Methods

### Simulation of anomalous diffusion using fractional Brownian motion

We simulated 2D single particle trajectories using fractional Brownian motion (fBM) using the ‘fbm_sim’ Python package (https://github.com/alecheckert/fbm_sim) (17) with anomalous exponents *α* = 0.2, 0.6, and 1 (Hurst coefficient H = *α /2*), and corresponding generalized diffusion coefficients D = 0.002, 0.025, and 0.8 µm^2^/s^*α*^, respectively. For each condition, 5000 trajectories of lengths 6, 10, 15, and 30 displacements were generated with a temporal spacing of 10ms. Zero-mean Gaussian noise with standard deviation *σ*_loc_ was added independently to the x and y coordinates of each trajectory point.

### Jump distance distribution analysis (JDD)

Custom Python code was used to fit jump distance histograms (bin width 2nm) with a fit equation describing a multi-component free diffusion probability distribution (24) :

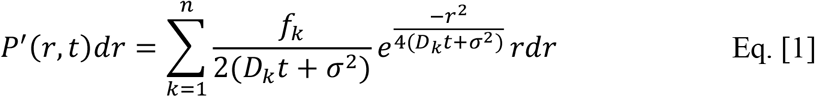

where (*f*_*k*_, *D*_*k*_) are the relative fraction and (Brownian) diffusion coefficient parameterizing specific mobility states, and *σ* the localization precision. The experimental localization precision can be estimated from independent experiments, e.g. imaging under the same conditions in fixed cells. Hence, we fixed *σ* to the input parameter for JDD analysis. To determine the number of mobility states in a track ensemble, we varied the number of assumed states *n* in the analysis from 1 to 5 and calculated the Bayesian information criterion (BIC) (25):

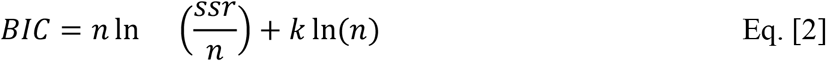

where ‘*ssr*’ is the sum of the square residuals, n is the number of observations, and k is the number of free parameters in the system (e.g., k = 5 for a three-state model, three diffusion coefficients and two independent relative fractions). The minimum BIC score informs on the appropriate model complexity.

### Bulk mean square displacement analysis (MSD)

Bulk mean-squared displacement (MSD) analysis was performed by fitting all data to the power law equation

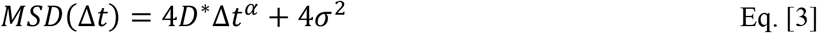

where *D*^*^ is the generalized diffusion coefficient, and *α* the anomalous exponent. Finite localization precision leads to an offset 4*σ*^2^ (26).

### Single-track mean square displacement analysis (stMSD)

For single-track MSD analysis, the mean square displacement as a function of lag time was calculated from coordinates in individual trajectories using custom python code. To minimize the number of free parameters, we omitted localization precision in the fit equation:

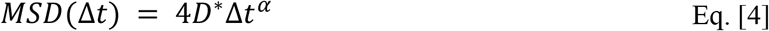

We fitted the first four data points (up to Δ*t* = 40*ms*) for each track with a minimum length of 6 coordinates (5 displacements) and plotted the distribution of parameters (*D*^*^, *α*) from single tracks in a scatter plot of *α*-vs-log(*D*^*^).

### Gaussian mixture model analysis

The resulting distribution of parameters (*D*^*^, *α*) was analyzed using a Gaussian mixture model (GMM) implemented in scikit-learn (27). We varied the number of components and used Bayesian information criterion scores to determine the justified number of GMM components (n_components). The model returns centroid coordinates in *α*-log(D^*^) space that we interpret as sub-population diffusion parameters (*D*^*^, *α*), relative fractions of each sub-population in the model via the mixture weights associated with each Gaussian component, and a covariance matrix describing the spread of parameters within each component.

Each Gaussian component is described by a probability density function

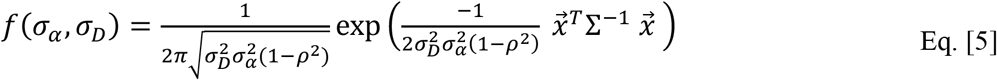

where 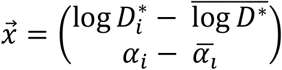 the centroid coordinates and 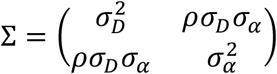 the covariance matrix with correlation *ρ* and standard deviations *σ*_*D*_ along log *D*^*^ and *σ*_*α*_ along the *α*-axis, respectively. The distribution of all datapoints in *α*-vs-log *D*^*^ space is then described by a linear combination of three such distributions with relative weights reflecting the proportion of tracks attributed to each component.

### Number of components in GMM analysis

We varied the number of components in Gaussian mixture model analysis using the Scikit-learn library (20) GaussianMixture estimator from 1 to 5 for mixed-ensemble data. The BIC score was computed using a standard implementation in GaussianMixture and used for model selection.

The optimal model size corresponds to the minimum BIC score.

### Odds ratio

For each trajectory, we obtained the probability of being in each state from GMM analysis. The odds ratio was calculated as the first highest to the second-highest probability amplitude for each track. Odds ratios close to 1 indicate greater uncertainty between the states. To remove ambiguous tracks, we rejected those with odds ratio ≤ 5.

### State array Single Particle tracking (saSPT)

We used publicly available code for FBME-based state array single particle tracking (saSPT) as described in the original publication (17). Diffusion coefficients were discretized over a logarithmically spaced grid ranging from 10^-3^ to 10^1^ µm^2^/s using 100 support points, while localization errors ranged from 0 to 450nm in 5nm increments. Hurst parameters *H* = *α*/2 ranged from 0.005 to 0.655 in increments of 0.05. We converted the saSPT output to an ensemble-wide posterior probability distribution in *α*-log(D^*^) space. While the original saSPT publication described using minima in log(D^*^) to separate states and assign fractions, we find this not feasible because posterior distributions overlap along this dimension. Sub-populations overlap more clearly in the posterior probability distribution of *α*. We visually determined diagonal lines that separate sub-populations in the posterior distribution and calculated fractions as the sum of posterior probabilities in each sub-region. Sub-population diffusion parameters (*D*^*^, *α*) were calculated as the weighted mean within the respective sub-regions in *α*-log(D^*^) space.

Internally, saSPT calculates a posterior distribution for each individual track. For single-track state assignment, we calculated parameters (*D*^*^, *α*) as the weighted mean of posterior probabilities. We then placed the parameter pair for each track in α-log(D^*^) space as described above for stMSD analysis.

### Single-track state assignments

Each point in α-log(D^*^) space inherits different probability densities from each component in a Gaussian mixture model. In the simplest approach, we assigned each track to the state with the highest local probability density at its coordinates. To exclude ambiguous assignments, we calculated the ratio of highest to second highest probability density (odds ratio). This odds ratio determines how much the most likely state is favored over the next most likely one and was used for thresholding.

### Calculation of recall rate and precision

The recall ratio was calculated for each state as the fraction of tracks reassigned to their originally simulated state after classification. It was calculated as the number of tracks assigned back to that state divided by the total number of tracks originally belonging to that state.

The precision was calculated as the fraction of all tracks assigned to a particular state during analysis that had indeed been simulated with the parameters of the respective state.

## Results

### Experimental analysis of RNA Pol II mobility in mouse embryonic stem cell nuclei

We previously reported analysis of an experimental dataset containing a total of 476,918 Snap-Pol II displacements in 128,563 tracks at Δ*t* = 10*ms* temporal resolution from 269 cells (18). The full dataset is publicly available (28). Sparse activation of a photo-activatable Snap-ligand (PA-JF646) allowed us to use a simple Hungarian algorithm with maximum jump length 600 nm to connect localizations to tracks (**Fig. 1a**).

**Fig. 1:**
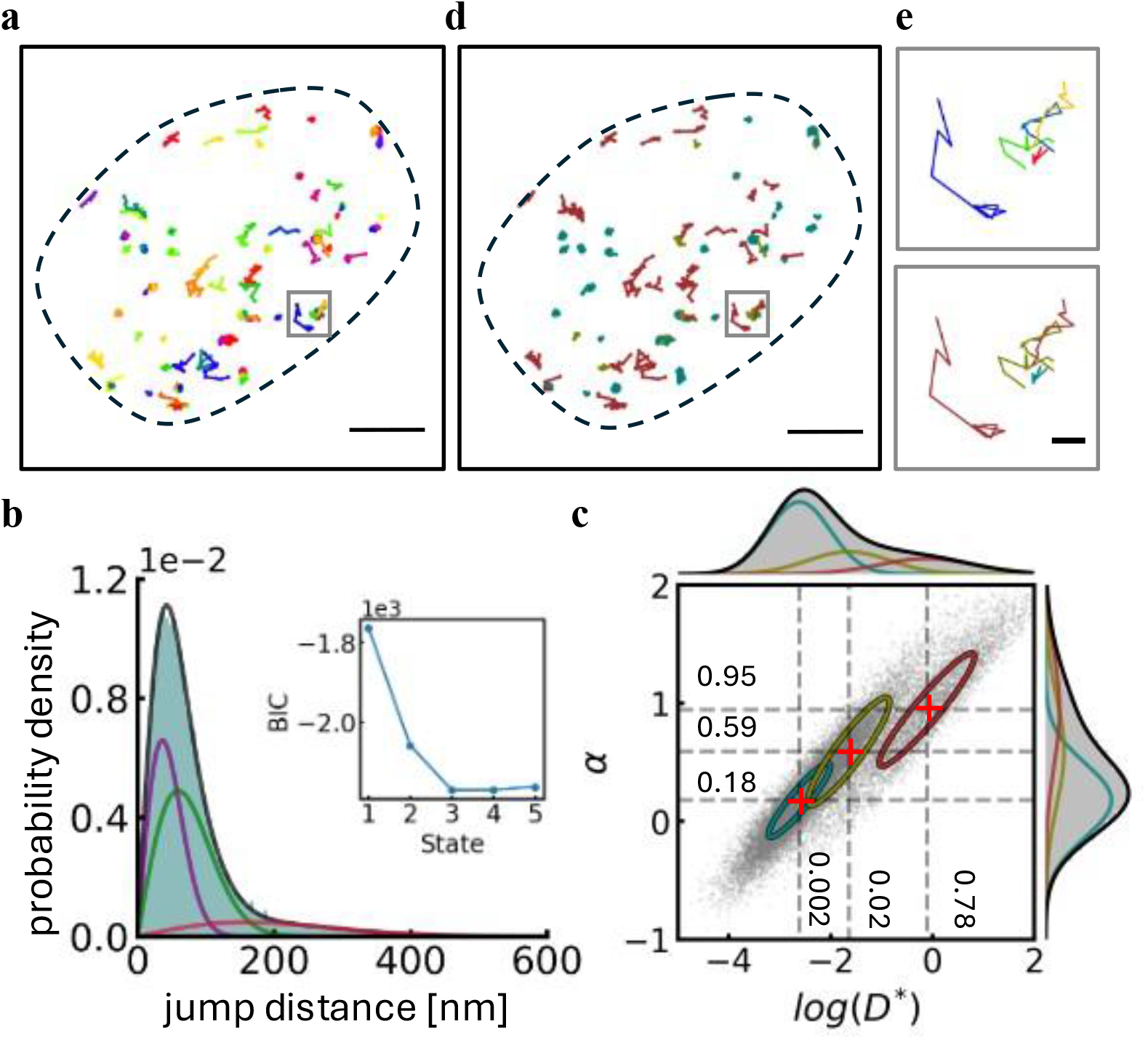
Analysis of experimental Pol II tracks. **a**) Pol II tracks from a single representative nucleus (black dashed line). Tracks are color-coded by track ID. **b**) Jump distance distribution analysis of tracks with minimum length 6 and three-state fit. Inset: BIC scores for different numbers of states in the analysis. **c**) α-vs-log(D*) plot obtained from single-track MSD analysis of the same data. The distribution is best described by a three-state model according to Gaussian mixture model analysis. **d**) Pol II tracks color-coded by mobility state determined from stMSD analysis. **e**) Magnified view of the area highlighted in **a**) with tracks color-coded by track ID (top) or mobility state (bottom). Scale bar: **a, d**) 2 µm, **e**) 250nm. Data published in (18) and available at figshare (28). This figure contains an analysis of tracks within the dataset of minimum length 6.

Jump distance distribution (JDD) analysis of all displacements with a free Brownian motion model yields a three-state model as the best fit according to Bayesian Information Criterion analysis (**Table 1**). We included an experimentally determined localization precision of *σ* = 22*nm* in the analysis (Eq. [1]). Limiting the analysis to a subset of 28,267 tracks with minimum length 5 displacements, i.e. at least 6 localizations, resulted in small shifts towards lower diffusion coefficients (**Fig. 1b, Table 1**). For the remainder of this manuscript, we refer to track length in terms of number of localizations. As expected, the highest mobility state *D*_3_ is underreported in the truncated dataset since fast-moving particles tend to diffuse out of the detection volume more quickly and are more susceptible to missed localization due to motion blur than lower mobility particles (14).

**Table 1:** Parametric descriptors of experimental Pol II mobility states determined from JDD and stMSD analysis. Mean +/- standard deviation from 50 iterations of fits to random samples (with replacement) of 25,000 jumps. Generalized diffusion coefficients for single-track MSD analysis have units of µm^2^/s^α^.

|  | state 1 |  | state 2 |  | state 3 |  |
| --- | --- | --- | --- | --- | --- | --- |
| | $f_1$ | $D_1 (\mu m^2/s^\alpha)$ | $f_2$ | $D_2 (\mu m^2/s^\alpha)$ | $f_3$ | $D_3 (\mu m^2/s^\alpha)$ |
| JDD<br>all tracks | $38 \pm 6\%$ | $0.02 \pm 0.01$ | $46 \pm 5\%$ | $0.17 \pm 0.02$ | $17 \pm 1\%$ | $1.5 \pm 0.1$ |
| JDD<br>length $\geq 6$ | $40 \pm 11\%$ | $0.02 \pm 0.01$ | $47 \pm 1\%$ | $0.13 \pm 0.03$ | $13 \pm 1\%$ | $1.2 \pm 0.1$ |
| stMSD<br>length $\geq 6$ | 57% | 0.002<br>($\alpha_1 = 0.18$) | 24% | 0.02<br>($\alpha_2 = 0.59$) | 19% | 0.78<br>($\alpha_3 = 0.95$) |

We also conducted a single-track mean-square displacement (stMSD) analysis on the subset of tracks with length ≥ 6. In this approach, each individual track is described by an anomalous exponent *α* and a generalized diffusion coefficient D^*^ by fitting its MSD with Eq. [4]. We subjected the resulting data distribution to Gaussian mixture model analysis (GMM) and used the Bayesian information criterion to determine a three-state model to be the best parameterized description of the full dataset (**Fig. 1c**). stMSD yields two low mobility states with sub-diffusive exponents (*α*_1_ = 0.18, *α*_2_ = 0.59), and one freely diffusive state (*α*_3_ = 0.95). stMSD analysis also allowed us to assign each track to one of the three mobility states and map these states back to the nuclear geometry (**Fig. 1d,e**), emphasizing the additional spatial information obtained from single track analysis.

Since stMSD data points result from fits to the anomalous diffusion equation (Eq. [4]) whereas JDD analysis assumes free diffusion (*α* = 1 for all states), parametric descriptions of the three states differ. stMSD analysis can yield free Brownian motion with an anomalous exponent *α* = 1 when appropriate, but JDD analysis cannot inform on anomalous behavior. The difference in models and scaling of displacements with lag time impacts (generalized) diffusion coefficients returned by the analyses. Further, each track contributes equally to relative fractions in stMSD analysis. Thus, fractions correspond to fractions of tracks or molecules in the respective state. In JDD analysis, the fractions report the number of jumps in each state (5,9). Hence, longer tracks, i.e. typically those of lower mobility, are overrepresented. Lastly, the jump distance distribution model (Eq. [1]) included a term for the experimental localization precision, whereas we omitted this offset in stMSD analysis to limit the number of free fit parameters when analyzing short tracks.

Even though we can rationalize the differences between the two analysis approaches, the question arises how datasets should be analyzed to obtain the best results, i.e. the most accurate parametric description of relative fraction, diffusion coefficient, and anomalous exponents for the sub-populations in an ensemble of tracks from different mobility states. Answering this question requires knowledge of the ground truth, which is not typically available for experimental data. Hence, we created *in silico* datasets by fractional Brownian motion simulation inspired by the experimental parameters determined by stMSD analysis (**Table 1**) to assess the accuracy of different analysis approaches.

### Bulk and single-track MSD analysis of single-component ensembles accurately reproduce anomalous diffusion input parameters

We simulated tracks using a fractional Brownian motion model suitable for modeling anomalous diffusion (17) to generate tracks with known ground truth as described above. We generated ensembles consisting of 5000 tracks of a fixed length (6, 10, 15, and 30 positions each) using a single set of anomalous diffusion input parameters, *D*^*^ and *α*. The combination of these two parameters determines a mobility “state”. To first ascertain the accuracy of the simulations, we ran 10 iterations and analyzed the track ensembles using JDD (free diffusion model), bulk MSD, and stMSD, as well as state-array single particle tracking (saSPT) analysis (17). We conducted this analysis for three mobility states individually with sets of parameters mirroring our experimental analysis results.

Results for track length 30 are illustrated in **Fig. 2**. All results are summarized in **Table 2**. For each of the three single-state ensembles, bulk MSD analysis (**Fig. 2b**) yields highly accurate results for both parameters, *D*^*^ and *α*, with little variation between iterations. As expected, the Brownian motion JDD analysis (**Fig. 2c**) fares equally well for the freely diffusive state 3.

**Table 2:** Input parameters for fractional Brownian motion simulations and values recovered from analysis of single-state ensembles with different methods. 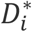 in units of 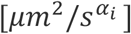. Values indicate mean ± s.d from 10 iterations.

|  | length | state 1 |  | state 2 |  | state 3 |  |
| --- | --- | --- | --- | --- | --- | --- | --- |
| | | $D_1^*$ | $\alpha_1$ | $D_2^*$ | $\alpha_2$ | $D_3^*$ | $\alpha_3$ |
| input |  | **0.002** | **0.20** | **0.025** | **0.60** | **0.80** | **1.00** |
| JDD | 10 | 0.0797<br>$\pm 0.0006$ | - | 0.157<br>$\pm 0.001$ | - | 0.800<br>$\pm 0.005$ | - |
| | 30 | 0.0796<br>$\pm 0.0002$ | - | 0.158<br>$\pm 0.001$ | - | 0.800<br>$\pm 0.002$ | - |
| bulk MSD | 10 | 0.0019<br>$\pm 0.0003$ | 0.22<br>$\pm 0.04$ | 0.025<br>$\pm 0.001$ | 0.61<br>$\pm 0.02$ | 0.81<br>$\pm 0.04$ | 1.01<br>$\pm 0.01$ |
| | 30 | 0.00194<br>$\pm 0.00006$ | 0.22<br>$\pm 0.02$ | 0.0250<br>$\pm 0.0008$ | 0.61<br>$\pm 0.02$ | 0.81<br>$\pm 0.02$ | 1.00<br>$\pm 0.01$ |
| stMSD | 10 | 0.00182<br>$\pm 0.00001$ | 0.194<br>$\pm 0.002$ | 0.0198<br>$\pm 0.0006$ | 0.563<br>$\pm 0.007$ | 0.52<br>$\pm 0.01$ | 0.918<br>$\pm 0.005$ |
| | 30 | 0.00194<br>$\pm 0.00002$ | 0.1985<br>$\pm 0.0003$ | 0.0234<br>$\pm 0.0002$ | 0.590<br>$\pm 0.002$ | 0.713<br>$\pm 0.001$ | 0.978<br>$\pm 0.003$ |
| saSPT | 10 | 0.069<br>$\pm 0.001$ | 0.32<br>$\pm 0.01$ | 0.143<br>$\pm 0.001$ | 0.646<br>$\pm 0.008$ | 0.770<br>$\pm 0.006$ | 1.040<br>$\pm 0.006$ |
| | 30 | 0.145<br>$\pm 0.001$ | 0.605<br>$\pm 0.004$ | 0.146<br>$\pm 0.001$ | 0.608±<br>0.004 | 0.787<br>$\pm 0.003$ | 1.012<br>$\pm 0.003$ |

**Fig. 2:**
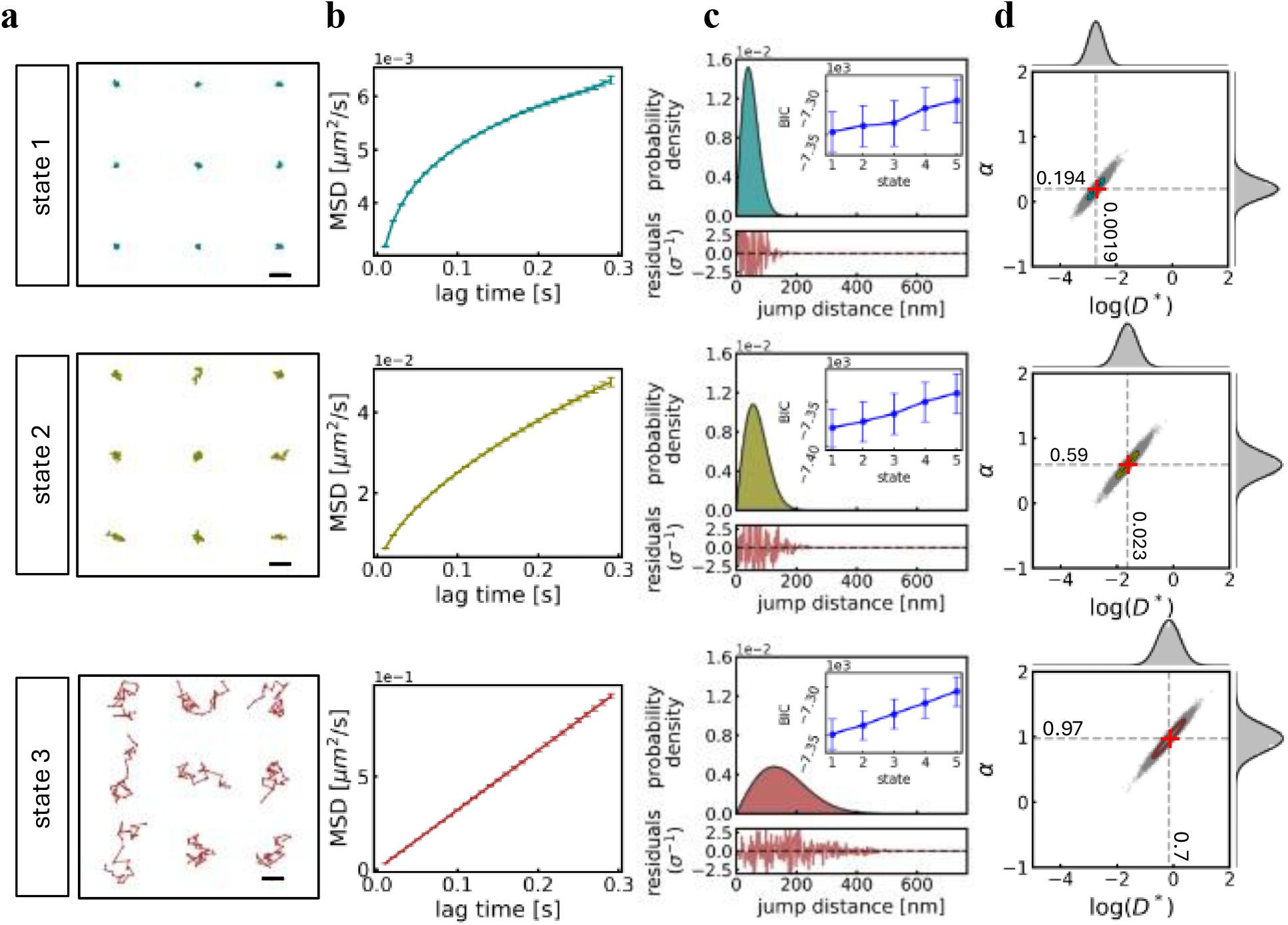
Analysis of simulated single-state anomalous diffusion ensembles. Simulations consisted of 5000 tracks each. Each of three states was simulated and analyzed independently. See **Table 2** for input parameters and output values. **a**) Example trajectories of length 30 coordinates at 10ms time resolution. **b**) Fits to bulk MSD data with an anomalous diffusion model. **c**) JDD analysis with a single component Brownian motion model (inset: BIC scores for 1-5 state models), that cannot inform on anomalous parameters. **d**) GMM analysis of the *α*-vs-log(D^*^) histogram obtained from stMSD. Bulk and stMSD analyses accurately recover input parameters. Scale bar in **a**), 500nm.

However, the sub-diffusive states 1 and 2 yield poor results for the diffusion coefficients (**Table 2**) since anomalous diffusion data were fit with a free diffusion model. Diffusion coefficients of the sub-diffusive states are overestimated. Notably, it is not possible to diagnose this issue from the JDD analysis itself. Criteria like BIC analysis or inspection of residuals indicate good single-state fits (**Fig. 2c**).

The stMSD analysis approach yields a spread of output parameters from single tracks (**Fig. 2d**). It is well-known that MSD analysis of single tracks is highly inaccurate and noisy (26,29). In our experimental and synthetic datasets, output parameters from single tracks in a given state present as elliptical distributions in *α*-vs-log(D^*^) space elongated along the diagonal, indicating correlation between the two parameters. The spread of data within each state mostly depends on track length, but like bulk MSD, population centroids determined by GMM analysis closely match input parameters (**Table 2**). This outcome is true for all tested track lengths from 10 – 30 positions. For track length 6, GMM analysis of stMSD data indicated a two-state model as best fit.

Finally, we analyzed the datasets with saSPT, a method that is suited to extract anomalous diffusion parameters from track ensembles (17). We find that saSPT accurately recovers the anomalous exponents of all three tested states (**Table 2**). It also yields an accurate estimate of the diffusion coefficient of the freely diffusive state 3, but overestimates the generalized diffusion coefficient of the low mobility sub-diffusive states. The output deviates more strongly from the input parameter for lower mobility tracks as discussed in the original publication.

This initial assessment confirms that the fractional Brownian motion simulations yield single-state track ensembles. JDD analysis is limited to Brownian components and hence fails to accurately recover parameters of anomalous diffusion populations. saSPT excels at recovering the anomalous exponents but has shortcomings in the description of low mobility states. Input parameters can be recovered accurately with both bulk MSD and stMSD analysis across a range of values.

### Parameter extraction from a multi-state population with limited track length and localization error

Experimental single-particle tracking datasets are typically more complex than the simple single-state ensembles characterized in **Fig. 2**. Photobleaching and axial particle motion limit track length, localizations contain measurement uncertainties determined by photon shot noise, background and readout noise, optical imperfections, and mechanical vibrations as well as motion blur (30). Practically, all of these effects contribute to finite localization precision.

Finally, experimental data usually comprises multiple mobility states.

We repeated the analyses presented above on datasets that mimic these conditions. First, we varied track length from 6 – 30 positions. Second, we included localization uncertainty by transforming each individual coordinate (*x*_*i*_, *y*_*i*_) within a track to (*x*_*i*_ + Δ*x, y*_*i*_ + Δ*y*), where (Δ*x*, Δ*y*) represent offsets drawn from a 2D normal distribution with spread *σ*_*xy*_ corresponding to the localization precision. In addition to the error-free tracks discussed above, we simulated ensembles with *σ*_*xy*_ = 20*nm*, close to our experimentally determined value, and at a reduced localization precision of *σ*_*xy*_ = 40*nm*. Finally, we pooled equal numbers of tracks from the three single-state datasets discussed above (5000 tracks each) to create mixed mobility state ensembles. For each combination of track length and localization precision, we generated ten different ensembles by randomly selecting tracks.

We then compared the performance of stMSD and saSPT analysis in recovering both diffusion parameters and relative fractions. Knowing that bulk MSD analysis is not suited for multi-component analysis, we omitted this method. We included JDD analysis to illustrate that just like for the single-state analysis (**Fig. 2**) it yields outputs in a free diffusion framework that would not raise suspicion of inaccuracy without further investigation.

In the analysis, we assumed knowledge of an experimentally defined localization precision, which can be estimated from, e.g., fixed cell experiments. Experimental data can contain varying degrees of localization precisions since motion blur affects highly diffusive states more than low mobility or bound states (17) unless experimental procedures are adjusted, e.g. by stroboscopic illumination (14). To facilitate comparison, we omitted this detail in this study.

The first task in the analysis workflow is to determine the number of states that should be included in any model. We did not specify the number of expected states *a priori*, but used criteria driven by the data to determine the most appropriate number in an unbiased way. For JDD and stMSD we calculated BIC scores. For saSPT we resorted to a more qualitative approach, separating states by manually segmenting the posterior probability state distribution approximately along minima between subpopulations.

JDD analysis of the mixed ensemble indeed yields a three-state model according to BIC scores as the best fit for short (**Fig. 3b**) and long tracks (**Fig. 3d**) despite the fact that two states in the ensemble are sub-diffusive. Relative fractions are reported accurately, and while diffusion coefficients do not match anomalous input parameters (**Table 3**) they do agree with the values obtained from the single-state analyses (**Table 2**).

**Table 3:**
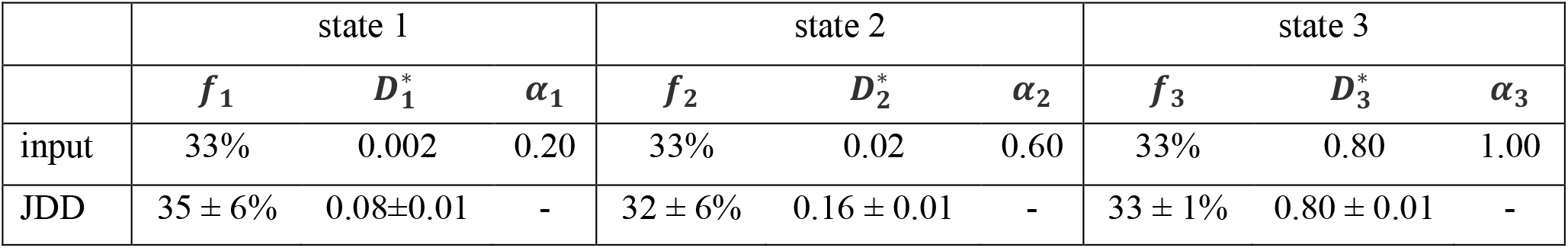
Parameters for fractional Brownian motion simulations and analysis recovered from analysis of a mixed population with JDD analysis.

|  | state 1 |  |  | state 2 |  |  | state 3 |  |  |
| --- | --- | --- | --- | --- | --- | --- | --- | --- | --- |
| | $f_1$ | $D_1^*$ | $\alpha_1$ | $f_2$ | $D_2^*$ | $\alpha_2$ | $f_3$ | $D_3^*$ | $\alpha_3$ |
| input | 33% | 0.002 | 0.20 | 33% | 0.02 | 0.60 | 33% | 0.80 | 1.00 |
| JDD | $35 \pm 6\%$ | $0.08 \pm 0.01$ | - | $32 \pm 6\%$ | $0.16 \pm 0.01$ | - | $33 \pm 1\%$ | $0.80 \pm 0.01$ | - |

**Fig. 3:**
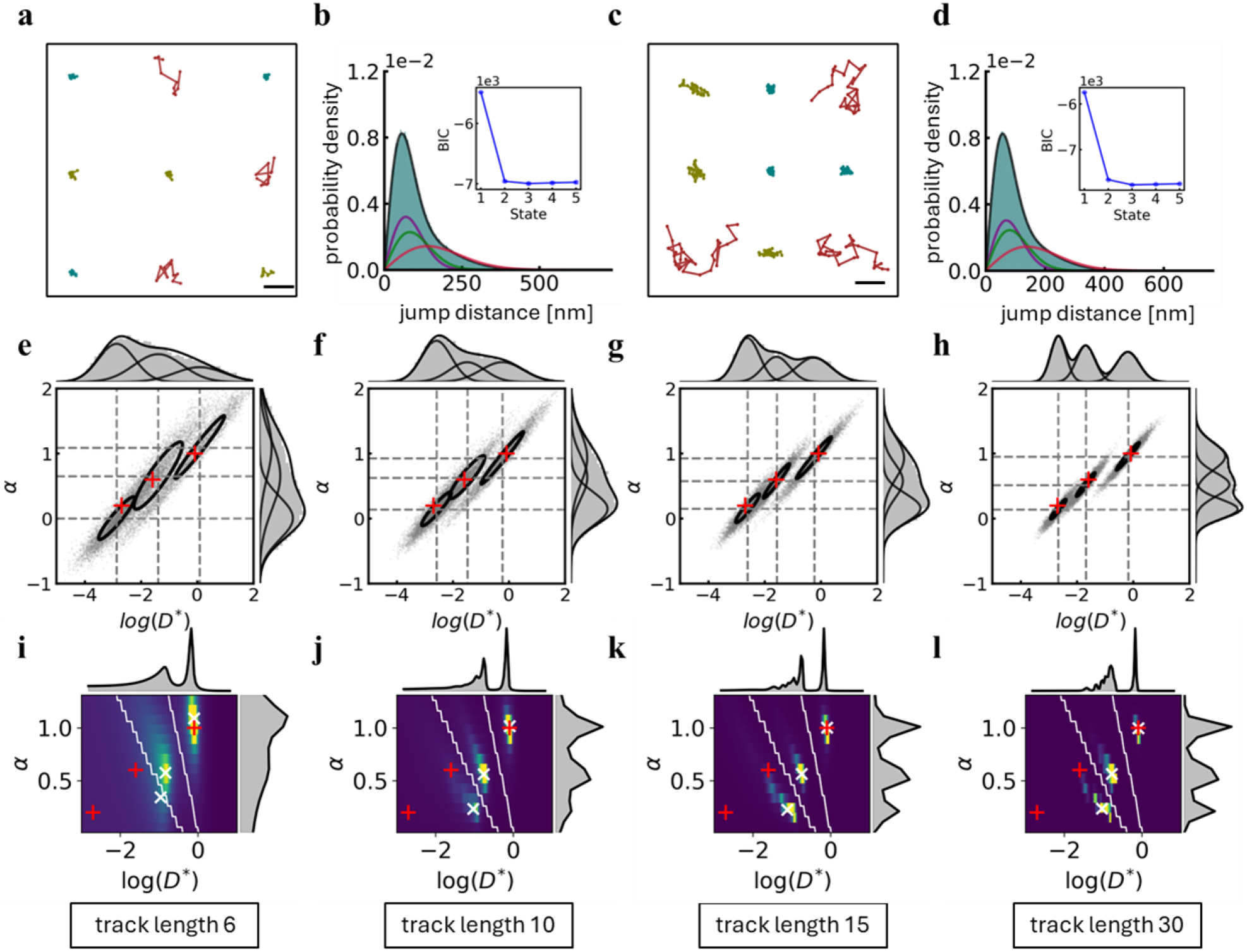
Mixed ensemble analysis with anomalous diffusion methods. We generated mixed ensembles with 5000 tracks from each of the three simulated components with localization precision of 20nm and varying track length (6-30). The simulation was repeated ten times for each condition. **a**) Example tracks with length 10. **b**) JDD analysis of the mixed ensemble yields three diffusive states. Inset: BIC scores for 1-5 state analysis. **c**) Example tracks with length 30. **d**) Corresponding JDD analysis. **e-h**) GMM analysis of the *α*-vs-log(D^*^) plots obtained from stMSD analysis for track length 6, 10, 15, and 30, respectively. Red crosses represent simulation input parameters. Gray dashed lines represent population centers recovered from the analysis. Black ellipses correspond to population spread determined from covariance matrices. Shorter tracks lead to larger spread of single-track output parameters but centroids are preserved. For track length 6, only 2/10 iterations returned a 3-state model as best fit (shown here). **i-l**) Bulk saSPT analysis of the same ensembles converted to posterior probability distributions in *α*-vs-log(D^*^) space. Yellow colors represent high posterior probability state occupancy. White x denotes population centers. White lines are drawn manually along minima in the distribution to quantify relative fractions. For track length 6, only two components were identified with systematic approaches in all iterations. Here, state boundaries from another track length were used to facilitate comparison. saSPT returns more distinct population centers, but the centroids of the lower-mobility components deviate strongly from the input parameters. Scale bar: **a**), **c**) 500nm.

In stMSD analysis, data from the different mobility states overlap in multi-component ensemble analysis (**Fig. 3e-g**). Shorter track length leads to a larger spread and greater overlap between states. Nevertheless, BIC assessment of GMM analysis recovers three mobility states reliably for tracks as short as 10 positions. For the shortest tracks (length 6), the number of states identified varied across iterations between three (2/10), four (7/10), and five (1/10). Simulations here included 15,000 tracks (5,000 from each state). It may be possible to improve the outcomes by including more tracks in the analysis (not tested here). The relative fractions assigned to the three states deviate from the input value by up to 15% for shorter tracks and low localization precision but are accurately reported for longer tracks under all simulated localization precision values. Centroids of sub-populations in the three-state Gaussian mixture model reproduce input parameters with good accuracy despite the increasing spread of each state (**Fig. 4a**). Generalized diffusion coefficients of all three states deviate from the input parameters only minimally. The anomalous exponent was also recovered well for most conditions with some deviations for the shortest track length. Here, each iteration of the simulation was analyzed with a three-state model for comparability, even though for track length 6 the majority of analysis runs on stMSD output returned four-state models as the best fit. All other track lengths resulted in three-state models. Inaccuracies for short tracks usually emerge when best fit covariance matrices in GMM analysis rotate towards a more vertical orientation. We noted above that the *α*-vs-log(D*) values of tracks in a single state exhibit a stereotypic correlation. This holds true across all simulation parameter combinations tested here. We speculate that constraining the orientation of the covariance matrix may alleviate issues with overlapping populations and improve outcomes.

**Fig. 4:**
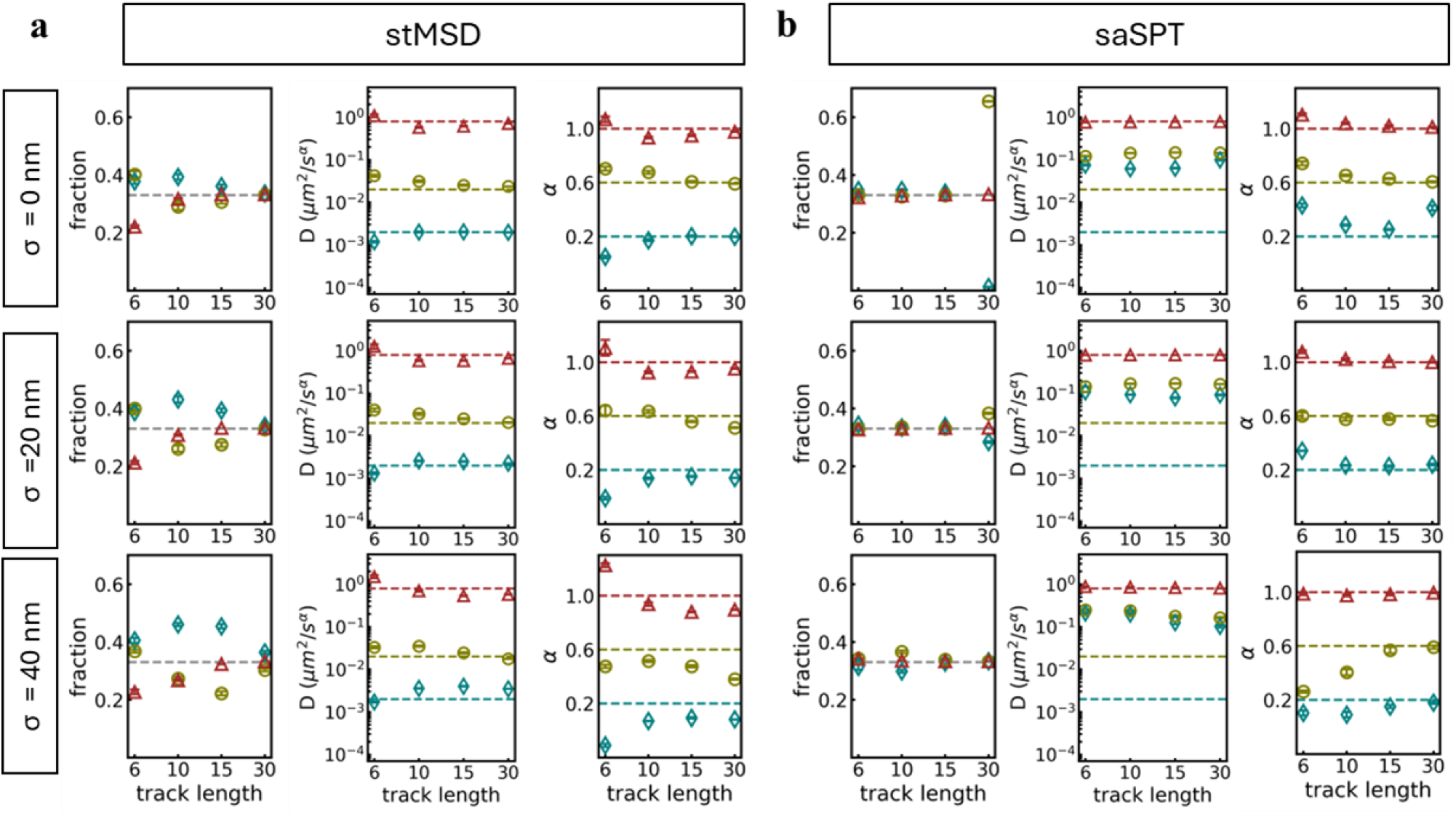
stMSD and saSPT result robustness for varying track length and localization precision. Mobility states in a mixed ensemble are defined by their relative fraction (left), generalized diffusion coefficient (middle), and anomalous exponent (right). We compare output parameters determined by **a**) stMSD and **b**) saSPT for localization precision of 0 nm (top), 20nm (middle), 40nm (bottom), and varying track length (6, 10, 15, and 30). Dashed lines represent ground truth input parameters. Data points for state 1 (teal), state 2 (yellow), and state 3 (red) are mean values ± s.d. from independent 10 iterations for each condition. For track length 6, both methods failed to reliably identify a three-state model as most appropriate. For comparability, we enforced a three-state description.

We also used saSPT analysis to assess the same mixed population ensembles (**Fig. 3i-l**). Generally, saSPT returned sharper peaks in the posterior distribution converted to *α*-vs-log (D*) space. The states are hence more readily distinguishable than in the stMSD output. To quantify relative fractions, we drew dividing lines in the posterior distribution that approximately followed the minima in that space. This approach yields near perfect recovery of the relative fractions for many conditions (**Fig. 4b**). We note however, that only two states could be identified for the shortest tracks with this method in all iterations (**Fig. 3i**). To facilitate comparison, we used the state mask obtained for track length 30. Paradoxically, quantification of relative fractions becomes worse for longer tracks and lower localization error (**Fig. 4b**). We were not able to find a satisfying explanation for this effect.

Anomalous exponents of all three states are well represented for tracks of length 10 – 30, with a localization error of 20nm even though the shortest tracks produced no distinguishable peak in the posterior distribution for the lowest mobility state (**Fig. 3i**). However, even for long tracks, generalized diffusion coefficients are severely overestimated for low-mobility states for all conditions that were simulated. The difficulty of saSPT in recovering parameters for low mobility states was previously described (17) and attributed to the impact of motion blur on localization error, but in our hands it persists even at *σ* = 0*nm*. The limitation likely arises from the intrinsic difficulty of resolving highly sub-diffusive populations. The freely diffusive state is described well even for the shortest tracks.

While Fig. 3 represents outcomes for simulations with localization precision of 20nm, close to our experimentally determined value, similar observations hold for simulations with no localization error or a localization precision of only 40nm. Generally, degrading localization precision increases the spread of outputs in both stMSD and saSPT, but the effect is far outweighed by the influence of track length. This influences recommendations for how to set up SPT experiments as discussed below.

### Assigning mobility states to individual tracks

In the analyses so far, we have focused on the ability to recover parametric descriptions of sub-populations in mixed ensembles, i.e. the input parameters for the simulation. Often, this information is sufficient to draw conclusions, e.g. when assessing the global effect of perturbations or mutations on single molecule dynamics. In some scenarios, however, it is of interest to assign mobility states to individual tracks. This is for example the case if the abundance of sub-populations in small sub-cellular regions or co-dependencies on other factors or events are of interest. Track-specific information is also important for counting molecules in a specific state.

Since diffusive particle motion is a stochastic process, it is never possible to declare states of individual tracks with 100% certainty unless sub-populations are well-separated by a quantifiable parameter. The overlapping populations in single-track analysis data (**Fig. 3**) illustrate this problem. Nevertheless, one can use single track information in relation to the full ensemble distribution to determine which state a given track most likely represents.

Here, we determine how accurate such assignments are by assessing two metrics, the “recall” rate and “precision” in single-track analyses (stMSD and saSPT) of mixed ensembles with known ground truth. “Recall” is defined as the probability that a track simulated as part of a given mobility state was also assigned to that state during analysis. The “precision” describes what percentage of tracks assigned to a state in the analysis were truly simulated with the corresponding diffusion parameters. We note that outcomes naturally will depend on the degree of overlap between probability density functions and the relative abundance of the three sub-populations. Here, we considered equal numbers of tracks in each of three states and parameter sets motivated by our experimental data. For clarity, we limit the analysis to tracks with realistic localization precision (20 nm). **Fig. 5** summarizes results for track length 10, the shortest tracks for which three states were unambiguously identified by population analysis.

**Fig. 5:**
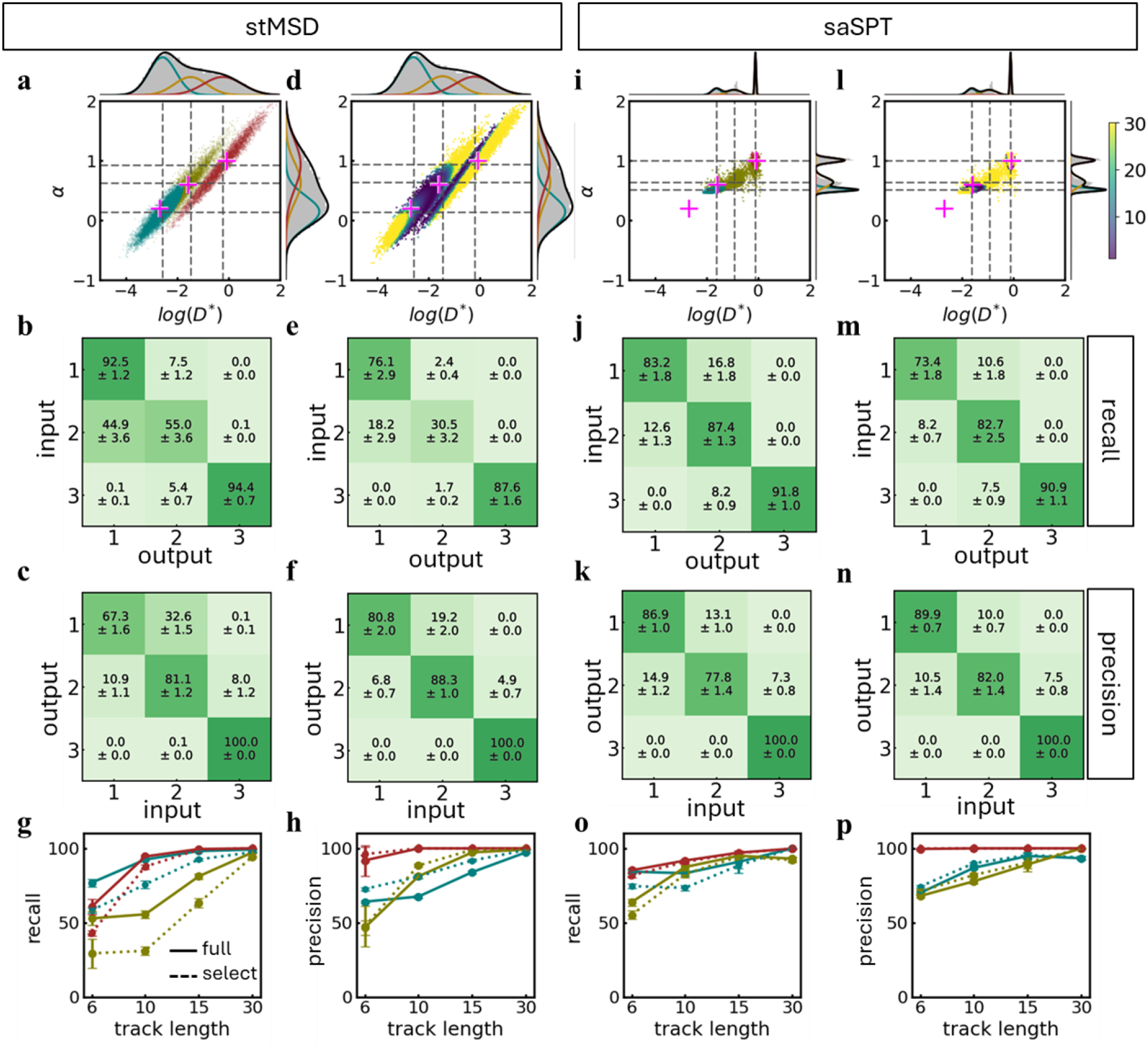
Single-track mobility state assignment. **a**) *α*-vs-log(D^*^) plot of stMSD analysis results for track length 10. Pink + indicates input parameters. Gray dashed lines population centers from GMM analysis. Each data point is color-coded by the state assigned to it by single-track analysis (teal: state 1, yellow: state 2, red: state 3). **b**) Recall matrix showing how likely tracks simulated in a specific state (input) are assigned to which output state. Diagonal entries represent recall fractions, i.e. the percentage of tracks simulated in a state that were assigned to the correct state. **c**) Precision matrix showing which input state the tracks assigned to each output state originated from. Diagonal entries are precision values, i.e. correctly identified tracks. **d**) Single-track output parameters color-coded by odds ratio, i.e. ratio of the highest to second highest probability density among the three states. We truncated ambiguous assignments by rejecting tracks with odds ratio ≤ 5. **e**) Recall values decrease because tracks predominantly from state 2 are rejected by thresholding. **f**) Instead, precision increases. **g**) Recall and **h**) precision for filtered and unfiltered data as a function of track length. **i**) saSPT output color-coded by single-track assignment. Pink + indicates input parameters. Gray dashed lines population centers. The spread of output data contracts, but population centers are more readily separated. **j**) Accordingly, the recall rate for state 2 and **k**) precision for state 1 improve compared to stMSD (**b**). **l**) Odds ratio thresholding of saSPT data has a small impact on **m**) recall and **n**) precision since the populations are better separated. **o**) Recall and **p**) precision of single-track assignment based on saSPT results as a function of track length.

For the assignment of states to individual tracks in stMSD and saSPT analysis, we made use of the outcomes of the GMM analysis. Besides centroids that define sub-population parametric descriptors, the GMM includes covariance matrices that define probability densities for each *α*-log(*D*^*^)-coordinate, i.e., for each specific track. In the simplest approach, we assigned each track the state with the highest local probability density (**Fig. 5a, c**). With this approach, stMSD yielded high recall values (>90%) for state 1 and 3 in a mixed ensemble of tracks with length 9 displacements (**Fig. 5b**). State 2 yielded low recall (55%) since it overlaps with state 1 to a high degree. Many state 2 tracks were erroneously assigned to state 1, but fewer state 1 tracks were assigned to state 2. Accordingly, the state 1 relative fraction was overrepresented (**Fig. 4a**). This effect also impacted precision, but to a lesser degree (**Fig. 5c**). Since 93% of tracks simulated to be in state 1 were also assigned to this state, the precision rate for state 1 remained at 67% despite misclassifications. Most tracks reported to be in state 1 indeed stemmed from this sub-population. Precision for state 2 (81%) and state 3 (100%) was even higher.

One strategy to improve precision is to exclude ambiguous assignments. To this end, we calculated the odds ratio of the highest to second-highest probability amplitude for each track (**Fig. 5d**). In a refined analysis, we considered tracks only if the odds ratio exceeded a minimum threshold of 5. As expected, excluding ambiguous assignments reduced recall rates. While 76.1% of all tracks passed the threshold, only 30.5% of tracks simulated in state 2 passed this criterion (**Fig. 5e**). However, as a result, precision improved to >80% for all three states (**Fig. 5f**).

We analyzed tracks of length 6-30 with both approaches (**Fig. 5g**,**h**). Thresholding improves precision most prominently for short tracks. Both recall and precision improve strongly for tracks of length 10 as compared to length 6 and approach perfect outcomes for the longest tracks (length 30) considered here.

We observed that saSPT analysis yielded more distinct populations in the posterior probability state distribution for most conditions (**Fig. 3i-l**). We reasoned that this might facilitate single track state assignments. As published, saSPT is a bulk analysis method, but in the background, posterior probability state distributions are calculated for each track. These distributions are accumulated to produce the outputs presented in **Fig. 3i-l**. We modified the approach by calculating a weighted average of the posterior state distribution for each single track. The outcome is a single pair of diffusive parameters per track similar to stMSD. We then plotted the value pairs in the same way (**Fig. 5i**). Since saSPT overestimated generalized diffusion coefficients of low mobility states, the overall distribution of output parameters is compressed to a smaller region in the scatter plot. The result is nevertheless a three-state distribution with clearly distinct peaks in *α*-log(*D*^*^)-space for track length 10-30. We then followed the same GMM analysis and single-track assignment procedure as described for stMSD.

Recall rates for all states are >80%, including for the intermediate state 2 that was most frequently mislabeled in stMSD. Recall rates for state 1 and 3 are slightly lower than for stMSD but still in a satisfactory range (**Fig. 5j**). Precision for state 1 increases for the same reason: Fewer state 2 tracks are assigned to state 1 (**Fig. 5k**). Precision for state 2 and 3 is comparable to stMSD output. Filtering by odds ratio (**Fig. 5l**) lowers the recall of state 1 (**Fig. 5m**) but has only a minor impact on the other states. This is not surprising given the fact that saSPT struggled most with the identification of the lowest mobility states (state 1) but also returned more focused output parameter distributions within each state, making it easier to determine clear state assignments. Accordingly, precision showed only minor improvements (**Fig. 5n**). Results improved further for longer tracks (**Fig. 5o,p**).

There are caveats to consider in single-track assignments: Firstly, stMSD and saSPT analysis failed to detect the correct number of states for the shortest tracks of length 6. It would not be meaningful to map two- or four-state analysis outcomes onto three-state input since there is no clear correspondence between input and output. To facilitate comparison, we enforced a three-state model in the analysis of short track ensembles. We note that experimental data consists of a distribution of track lengths. Length 6 was set as a lower limit here, and the results indicate that this was an appropriate choice since it is the regime where state identification begins to fail. Practically, the number of states could also be gleaned from comparative analysis with other methods, e.g. concurrent JDD analysis.

Further, even for longer tracks of length 15 and 30, parametric descriptors of low mobility states are highly overestimated by saSPT analysis. Hence, single tracks were confidently assigned to a state but with inaccurate parametric description. Whether or not this will be considered a problem depends on the goal of the analysis.

## Discussion

In this work, we used simulations to construct synthetic datasets comprising sub-populations of single particle tracks including anomalous diffusion behavior typically observed in the cell nucleus. We simulated tracks of varying length and with realistic degrees of localization precision. Classical methods for SPT data analysis are suited only for single-state scenarios (bulk MSD) or can return the number of states in the ensemble (JDD) but not report anomalous diffusion parameters. Analysis of experimental datasets thus requires more sophisticated methods to recover anomalous diffusion parameters.

We compared the ability of two methods, stMSD and saSPT to recover anomalous diffusion parameters. Both methods identified the correct number of states for track lengths as short as 10 localizations. saSPT returned accurate anomalous exponents but overestimated generalized diffusion coefficients even for the longest tracks under consideration. stMSD sub-populations broadened for shorter track length but population centroids remained true to the input values. We also used both methods to characterize recall and precision when assigning individual tracks to mobility states. When all three states were identified in the ensemble in saSPT, the single-track state assignments were more confident since states are more clearly separated in the posterior probability state distribution. stMSD success in this task was limited by the broad spread and resulting overlap of sub-populations in *α*-log(*D*^*^)-space. Precision in stMSD was improved by odds ratio thresholding, i.e. rejection of ambiguous data points, but at the cost of reduced recall. Thresholding was not necessary for saSPT, but with the caveat that diffusion parameters of low mobility states were severely overestimated. Such states are frequently present in tracking data, e.g. due to binding or sub-diffusive target search of single molecules. Accurate numerical values are important for biophysical interpretation.

Our results show that no single analysis framework is ideally suited to fully analyze a mixed ensemble of anomalous diffusion states. A prudent recommendation is to include results from multiple methods in the description of experimental tracking data. In a first step, the number of mobility states should be identified. Simple jump distance analysis can give an early indication but will yield only qualitative descriptions of non-Brownian states. In a second step, relative fractions and accurate anomalous diffusion parameters can be obtained. Here, saSPT and stMSD provide the advantage of yielding fractions in terms of the number of molecules rather than the number of displacements observed in a given state. We find that stMSD provides accurate diffusion parameters even for highly sub-diffusive states and short tracks, in particular if the number of states is known from an orthogonal analysis.

One notable realization emerging from our characterizations is that track length is more important than localization precision for both saSPT and stMSD analysis (within the ranges explored here). A consequence of this finding is that it may in fact be advisable to tolerate lower localization precision in favor of more observations of the same particle. With a finite photon budget reducing excitation power by a factor of two should in principle lower the bleaching rate by the same factor, i.e. double the average track length. This comes at the cost of a reduced localization precision *σ*. But since *σ* scales approximately as (*N*_*phot*_) ^−1/2^, localization precision would only degrade by a factor of 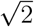. This tradeoff can be tolerated since the gain of longer tracks outweighs the cost of reduced localization precision.

Practical realities, such as fast diffusive particles leaving the detection range, of course set limits to this consideration. But low mobility states with highly sub-diffusive behavior that are most difficult to resolve by both methods tested here should be relatively robust to this modification and benefit the most. Additional measures such as stroboscopic illumination to reduce motion blur differences between low and high mobility particles are orthogonal and can provide further benefits.

The studies presented in this manuscript were motivated by a specific experimental scenario, the investigation of RNA Pol II single particle tracking data. Results from this experimental dataset served as a template for the specific values chosen for the simulated tracks. While concrete numerical values, e.g. error rates and uncertainties, are specific to the set of parameters, the fundamental conclusions of this work apply universally.

A limitation of our analysis is that we assume tracks to represent particles in a single state without intermittent transitions between states. This is justified especially for short tracks – tracks in our datasets correspond to 60-300ms lab time, at or below the times typically assigned to even short-lived “non-specific” interactions. Transitions within a single track would occur more frequently in longer tracks (16). Dedicated methods exist to dissect this type of data (12).

We believe that, though laborious, the comparative approach taken here provides a general template for scrutinizing experimental tracking data and quantifying uncertainties in the analysis. This may also involve additional tools (14,31) and approaches (lag time- and jump length-dependent angular anisotropy) (6) that we did not discuss here.

## Data availability

Experimental tracking data summarized in Fig. 1 that motivated the choice of simulation parameters is available through the [[Spille Lab FigShare repository](https://figshare.com/account/mycontent/projects/253574)]. Custom analysis code written in Python with usage instructions is available through the Spille Lab GitHub site [Spille-Lab/Pol-II-tracking-in-condensates: Code for single particle tracking and data analyses of Pol II mobility in transcription condensates.].

## Author contributions

Conceptualization: A.B., G.P., J.G-S., and J.H.S. Methodology: A.B., G.P., and J.H.S. Investigation: A.B. and J.H.S. Formal analysis: A.B. Software: A.B., and G.P. Visualization: A.B. Funding acquisition: J.H.S. Project administration: J.H.S. Supervision: J.H.S. Writing – original draft: A.B., and J.H.S. Writing – review & editing: A.B., G.P., J.G-S., and J.H.S.

## Acknowledgements

J.H.S. acknowledges support from the Research Corporation for Scientific Advancement (CMC 28407), NIH (R35GM150560), NSF (MCB-2306187) and the University of Illinois Chicago Startup Fund. This research was supported in part by grants from the NSF (DMS-2235451) and Simons Foundation (MP-TMPS-00005320) to the NSF-Simons National Institute for Theory and Mathematics in Biology (NITMB).

## References

1. Jinek M, Chylinski K, Fonfara I, Hauer M, Doudna JA, Charpentier E. A Programmable Dual-RNA–Guided DNA Endonuclease in Adaptive Bacterial Immunity. Science. 2012;337(C09C):81C–21. doi:10.112C/science.1225829

2. Zhang Y, Zheng Y, Tomassini A, Singh AK, Raymo FM. Photoactivatable Fluorophores for Bioimaging Applications. ACS Appl Opt Mater. 2023 Mar 24;1(3):C40–51. doi:10.1021/acsaom.3c00025 PubMed PMID: 37C01830; PubMed Central PMCID: PMC10437147.

3. Grimm JB, English BP, Chen J, Slaughter JP, Zhang Z, Revyakin A, et al. A general method to improve fluorophores for live-cell and single-molecule microscopy. Nature Methods. 2015 Mar;12(3):244–50. doi:10.1038/nmeth.325C

4. Grimm JB, English BP, Choi H, Muthusamy AK, Mehl BP, Dong P, et al. Bright photoactivatable fluorophores for single-molecule imaging. Nat Methods. 201C Dec;13(12):985–8. doi:10.1038/nmeth.4034

5. Hansen AS, Woringer M, Grimm JB, Lavis LD, Tjian R, Darzacq X. Robust modelbased analysis of single-particle tracking experiments with Spot-On. Sherratt D, editor. eLife. 2018 Jan;7:e33125. doi:10.7554/eLife.33125

6. Izeddin I, Récamier V, Bosanac L, Cissé II, Boudarene L, Dugast-Darzacq C, et al. Single-molecule tracking in live cells reveals distinct target-search strategies of transcription factors in the nucleus. Singer RH, editor. eLife. 2014 Jun 12;3:e02230. doi:10.7554/eLife.02230

7. Chen J, Zhang Z, Li L, Chen BC, Revyakin A, Hajj B, et al. Single-Molecule Dynamics of Enhanceosome Assembly in Embryonic Stem Cells. Cell. 2014 Mar;15C(C):1274–85. doi:10.101C/j.cell.2014.01.0C2

8. Kent S, Brown K, Yang C hsun, Alsaihati N, Tian C, Wang H, et al. Phase-Separated Transcriptional Condensates Accelerate Target-Search Process Revealed by Live-Cell Single-Molecule Imaging. Cell Reports. 2020 Oct 13;33(2):108248. doi:10.101C/j.celrep.2020.108248

9. Mazza D, Abernathy A, Golob N, Morisaki T, McNally JG. A benchmark for chromatin binding measurements in live cells. Nucleic Acids Research. 2012 Aug 1;40(15):e119–e119. doi:10.1093/nar/gks701

10. Ryosuke Nagashima. Single nucleosome imaging reveals loose genome chromatin networks via active RNA polymerase II. J Cell Biol. 2019;218(5):1511–30.

11. Wagh K, Stavreva DA, Jensen RAM, Paakinaho V, Fettweis G, Schiltz RL, et al. Dynamic switching of transcriptional regulators between two distinct low-mobility chromatin states. Science Advances. 2023 Jun 14;9(24):eade1122. doi:10.112C/sciadv.ade1122

12. Persson F, Lindén M, Unoson C, Elf J. Extracting intracellular diffusive states and transition rates from single-molecule tracking data. Nature Methods. 2013 Mar 1;10(3):2C5–9. doi:10.1038/nmeth.23C7

13. Gebhardt JCM. Single-molecule Tracking and Kinetic Analysis in Living Cells and Multicellular Organisms. Journal of Molecular Biology. 202C Jan;438(1):1C9308. doi:10.101C/j.jmb.2025.1C9308

14. Hansen AS, Woringer M, Grimm JB, Lavis LD, Tjian R, Darzacq X. Robust model-based analysis of single-particle tracking experiments with Spot-On. eLife Sciences. 2018 Jan 4;7:e33125. doi:10.7554/eLife.33125

15. Neumann FR, Dion V, Gehlen LR, Tsai-Pflugfelder M, Schmid R, Taddei A, et al. Targeted INO80 enhances subnuclear chromatin movement and ectopic homologous recombination. Genes Dev. 2012 Feb 15;2C(4):3C9–83. doi:10.1101/gad.17C15C.111

16. Spille JH, Kaminski TP, Scherer K, Rinne JS, Heckel A, Kubitscheck U. Direct observation of mobility state transitions in RNA trajectories by sensitive single molecule feedback tracking. Nucleic Acids Res. 2015 Jan;43(2):e14. doi:10.1093/nar/gku1194 PubMed PMID: 25414330; PubMed Central PMCID: PMC4333372.

17. Heckert A, Dahal L, Tjian R, Darzacq X. Recovering mixtures of fast-diffusing states from short single-particle trajectories. Smal I, Akhmanova A, Paul MW, editors. eLife. 2022 Sep C;11:e701C9. doi:10.7554/eLife.701C9

18. Spille JH, Budhathoki A, Pandey G, Medhanie F, Ngo GH, Mahmood S. Chromatin Binding Enriches RNA Polymerase II in Transcription Condensates [Internet]. Research Square; 2025 [cited 2025 Dec 15]. Available from: https://www.researchsquare.com/article/rs-7302C34/v1 doi:10.21203/rs.3.rs-7302C34/v1

19. Abrahamsson S, Chen J, Hajj B, Stallinga S, Katsov AY, Wisniewski J, et al. Fast multicolor 3D imaging using aberration-corrected multifocus microscopy. Nat Methods. 2013 Jan;10(1):C0–3. doi:10.1038/nmeth.2277

20. Boehning M, Dugast-Darzacq C, Rankovic M, Hansen AS, Yu T, Marie-Nelly H, et al. RNA polymerase II clustering through carboxy-terminal domain phase separation. Nat Struct Mol Biol. 2018 Sep;25(9):833–40. doi:10.1038/s41594-018-0112-y

21. Nguyen VQ, Ranjan A, Liu S, Tang X, Ling YH, Wisniewski J, et al. Spatiotemporal coordination of transcription preinitiation complex assembly in live cells. Molecular Cell. 2021 Sep 2;81(17):35C0–3575.eC. doi:10.101C/j.molcel.2021.07.022

22. Saxton MJ. Single-particle tracking: the distribution of diffusion coefficients. Biophysical Journal. 1997 Apr;72(4):1744–53. doi:10.101C/S000C-3495(97)78820-9

23. Pandey et. al. Characterizing Properties of Biomolecular Condensates Below the Diffraction Limit In Vivo. Phase-Separated Biomolecular Condensates: Methods and Protocols, Methods in Molecular Biology. 2023;25C3.

24. Siebrasse JP, Veith R, Dobay A, Leonhardt H, Daneholt B, Kubitscheck U. Discontinuous movement of mRNP particles in nucleoplasmic regions devoid of chromatin. Proceedings of the National Academy of Sciences. 2008 Dec 23;105(51):20291– C. doi:10.1073/pnas.0810C92105

25. Schwarz G. Estimating the Dimension of a Model. The Annals of Statistics. 1978 Mar;C(2):4C1–4. doi:10.1214/aos/117C34413C

26. Michalet X. Mean Square Displacement Analysis of Single-Particle Trajectories with Localization Error: Brownian Motion in Isotropic Medium. Phys Rev E Stat Nonlin Soft Matter Phys. 2010 Oct;82(4 Pt 1):041914. doi:10.1103/PhysRevE.82.041914 PubMed PMID: 21230320; PubMed Central PMCID: PMC3055791.

27. Pedregosa F, Varoquaux G, Gramfort A, Michel V, Thirion B, Grisel O, et al. Scikit-learn: Machine learning in Python. Journal of Machine Learning Research. 2011;12(Oct):2825–30.

28. Available at: https://figshare.com/articles/dataset/Pol_II_tracking_data_for_the_untreated_case_/29373998/1 doi:10.C084/M9.FIGSHARE.29373998.V1

29. Backlund MP, Joyner R, Moerner WE. Chromosomal locus tracking with proper accounting of static and dynamic errors. Phys Rev E. 2015 Jun 29;91(C):0C271C. doi:10.1103/PhysRevE.91.0C271C

30. Mortensen KI, Churchman LS, Spudich JA, Flyvbjerg H. Optimized localization analysis for single-molecule tracking and super-resolution microscopy. Nat Methods. 2010 May;7(5):377–81. doi:10.1038/nmeth.1447 PubMed PMID: 203C4147; PubMed Central PMCID: PMC3127582.

31. Martens KJA, Turkowyd B, Hohlbein J, Endesfelder U. Temporal analysis of relative distances (TARDIS) is a robust, parameter-free alternative to single-particle tracking. Nature Methods. 2024 Jun 1;21(6):1074–81. doi:10.1038/s41592-023-02149-7

32. Izeddin I, Récamier V, Bosanac L, Cissé II, Boudarene L, Dugast-Darzacq C, et al. Single-molecule tracking in live cells reveals distinct target-search strategies of transcription factors in the nucleus. eLife. 2014 Jun 12;3:e02230. doi:10.7554/eLife.02230

